# Perfluorooctanoic acid activates multiple nuclear receptor pathways and skews expression of genes regulating cholesterol homeostasis in liver of humanized PPARα mice fed an American diet

**DOI:** 10.1101/2020.01.30.926642

**Authors:** JJ Schlezinger, H Puckett, J Oliver, G Nielsen, W Heiger-Bernays, TF Webster

## Abstract

Humans are exposed to per- and polyfluoroalkyl substances (PFAS) in their drinking water, food, air, dust in their homes, and by direct use of consumer products. Increased concentrations of serum total cholesterol and low density lipoprotein cholesterol are among the endpoints best supported by epidemiology. The objectives of this study were to generate a new model for examining PFAS-induced dyslipidemia and to conduct molecular studies to better define mechanism(s) of action. We tested the hypothesis that PFOA exposure at a human-relevant level dysregulates expression of genes controlling cholesterol homeostasis in livers of mice expressing human PPARα (hPPARα). Female and male hPPARα and PPARα null mice were fed a diet based on the “What we eat in America” analysis and exposed to perfluorooctanoic acid (PFOA) in drinking water (8 µM) for 6 weeks. This resulted in a serum PFOA concentration of 48 μg/ml. PFOA increased liver mass, which was associated with histologically-evident lipid accumulation. PFOA induced PPARα and constitutive androstane receptor target gene expression in liver. Expression of genes in four pathways regulating cholesterol homeostasis were also measured. PFOA decreased expression of *Hmgcr* in a PPARα-dependent manner. PFOA decreased expression of *Ldlr* and *Cyp7a1* in a PPARα-independent manner. *Apob* expression was not changed. Gene expression in females appeared to be more sensitive to PFOA exposure than in males. This novel study design (hPPARα mice, American diet, long term exposure) generated new insight on the effects of PFOA on cholesterol regulation in the liver and the role of hPPARα.

## Introduction

Per- and polyfluoroalkyl substances (PFAS) are pervasive in the environment because of their persistence and extensive use in consumer products and fire-fighting foam. Daily human exposures occur via PFAS contaminated food, drinking water, dust and air (EFSA 2018; Kim et al. 2019; Makey et al. 2017). Multiple adverse health outcomes have been associated with PFAS exposure including birth outcomes, immunologic effects, and metabolic disruption. Among the best supported and most sensitive endpoints in both cross-sectional and longitudinal epidemiology studies are lipid-disrupting effects (Frisbee et al. 2010; Geiger et al. 2014; Graber et al. 2018; He et al. 2018; Nelson et al. 2010; Steenland et al. 2009). The European Food Safety Authority has proposed dyslipidemia, an abnormal amount of lipids (triglycerides, cholesterol and/or fat phospholipids) in the blood, as a critical effect (EFSA 2018).

Mechanisms by which PFAS may cause lipid-disrupting effects are not well understood. Fatty acids are endogenous ligands for peroxisome proliferator activated receptor α (PPARα), a transcription factor that regulates lipid homeostasis. Since many PFAS are structurally similar to fatty acids, PPARα binding and activation is a logical molecular initiating event for a lipid-disrupting pathway. As such, PPARα was found to account for 80-90% of perfluorooctanoic acid (PFOA) regulated genes (Rosen et al. 2017). There are well known species differences in the function of mouse PPARα (mPPARα) and human PPARα (hPPARα) (Gonzalez and Shah 2008). Activation of mPPARα results in peroxisome proliferation and dysregulation of cell cycle genes, which does not occur in humans (Morimura et al. 2006). However, both mPPARα and hPPARα efficaciously regulate multiple biological pathways involved in lipid homeostasis, acute phase response and inflammation (Rakhshandehroo et al. 2009). mPPARα and hPPARα share 91% amino acid identity (Sher et al. 1993), and important differences in the ligand binding domains have been identified (Keller et al. 1997; Oswal et al. 2014). As a result, there are differences in ligand specificity and gene expression patterns (Keller et al. 1997; Oswal et al. 2014; Rakhshandehroo et al. 2009). However, both mPPARα and hPPARα have been shown to contribute to regulation of cholesterol homeostasis (Bouchard-Mercier et al. 2011; Flavell et al. 2000; Peters et al. 1997; Robitaille et al. 2004; Sparso et al. 2007; Tanaka et al. 2007; Vohl et al. 2000).

Studies using reporter assays show that hPPARα is activated by PFAS. hPPARα activation increases with perfluoroalkyl acid chain length up to 8 carbons, yet all carbon chain lengths tested (i.e., C4 to C12) significantly activate PPARα (Behr et al. 2019; Maloney and Waxman 1999; Rosenmai et al. 2018; Takacs and Abbott 2007; Vanden Heuvel et al. 2006). hPPARα target gene expression is induced by PFAS in human hepatocyte models, including primary hepatocytes (Behr et al. 2019; Bjork et al. 2011; Buhrke et al. 2013; Buhrke et al. 2015; Peng et al. 2013; Rosen et al. 2013; Wolf et al. 2012; Wolf et al. 2008). However, in human and rodent hepatocyte models, transcriptome profiling shows that PFAS also regulate target gene expression of other nuclear receptors (Abe et al. 2017; Bjork et al. 2011; Buhrke et al. 2015; Scharmach et al. 2012).

Studies of the effects of PFOA on serum lipids and cholesterol in animal models have produced contradictory results. Mice exposed to PFOA and fed standard rodent chow (low in fat with negligible cholesterol) show decreased serum cholesterol levels (reviewed in (Rebholz et al. 2016)), in contrast to the increase shown in human epidemiology. But diet also influences serum cholesterol levels in mice (Dietschy et al. 1993), and when mice are fed a cholesterol and fat-containing diet, PFOA does induce hypercholesterolemia (Rebholz et al. 2016). Strain and sex also modify PFAS-induced effects on serum lipids (Pouwer et al. 2019; Rebholz et al. 2016).

Last, in mice expressing human PPARα in liver, PFOA increased serum cholesterol (Nakamura et al. 2009). The goals of this study were to generate a new model for examining PFAS-induced dyslipidemia and to conduct molecular studies to better define the mechanism of action by which this occurs. We tested the hypothesis that PFOA exposure dysregulates genes controlling cholesterol homeostasis in livers of mice expressing human PPARα (hPPARα) and fed an American diet, the first time this combination has been examined. We focused on liver because it is an essential site of regulation of multiple aspects of cholesterol homeostasis (Dietschy et al. 1993). The role of hPPARα was determined through comparison with effects in PPARα null mice. The data document important sex differences and identification of molecular pathways important for PFAS-induced dyslipidemia.

## Materials and Methods

### Materials

Perfluorooctanoic acid (cat. # 171468, 95% pure) was from Sigma-Aldrich (St. Louis, MO). All other reagents were from Thermo Fisher Scientific (Waltham, MA), unless noted.

### In vivo exposure

All animal studies were approved by the Institutional Animal Care and Use Committee at Boston University and performed in an American Association for the Accreditation of Laboratory Animal Care accredited facility (Animal Welfare Assurance Number: A3316-01). Male and female, humanized PPARα mice (hPPARα) were generated from mouse PPARα null, human PPARα heterozygous breeding pairs (generously provided by Dr. Frank Gonzalez, NCI)(Yang et al. 2008). Experiments were carried out using 11 cohorts of mice generated from four breeding pairs (Table S1). Genotyping for mouse and human PPARα was carried out by Transnetyx (Cordova, TN). The expression level of hPPARα in liver was confirmed by RT-qPCR.

At weaning, mice were provided a custom diet based on the “What we eat in America (NHANES 2013/2014)” analysis for what 2-19 year old children and adolescents eat (Research Diets, New Brunswick, NJ)(USDA 2018). The diet contains 51.8% carbohydrate, 33.5% fat, and 14.7% protein, as a % energy intake (Table S2). Fats are in the form of soybean oil, lard and butter, with cholesterol at 224 mg/1884 kcal. Vehicle (Vh) and treatment water were prepared from NERL High Purity water (23-249-589, Thermo Fisher Scientific), which is prepared using the most efficacious methods to remove PFAS (i.e., reverse osmosis and carbon filtering)(Appleman et al. 2014). A concentrated stock solution of PFOA (1×10^−2^ M) was prepared in NERL water and then diluted in NERL water containing 0.5% sucrose. Mice were administered vehicle (0.5% sucrose) drinking water or PFOA (8 μM) drinking water *ad libitum* for 6-7 weeks. Food and water consumption were determined on a per cage basis each week. Body weight was measured weekly. Mice were analyzed for body composition (total body fat mass, lean mass, water) using an EchoMRI700 (EchoMRI LLC, Houston, TX), fasted for 6 hours and then euthanized. Livers were collected from each mouse and weighed. Aliquots of liver for gene expression were flash frozen in liquid nitrogen and stored at −80°C and for histology were fixed in 10% neutral buffered formalin.

### PFAS analysis

PFAS concentrations were determined in pooled water samples. Aliquots were taken each time drinking water was made for a cohort and combined in a single sample; drinking water from five cohorts was analyzed. Individual serum samples were analyzed. PFOA concentrations were determined by LC-MS/MS according to methods MLA-110 (EPA Method 537 Modified) for water and MLA-042 for serum (SGS AXYS Analytical Services Ltd., Sidney, British Columbia, CA).

### Gene expression analyses

Total RNA was extracted and genomic DNA was removed using the Direct-zol RNA Miniprep Kit (Zymo Research, Orange, CA). cDNA was synthesized from total RNA using the iScript™ Reverse Transcription System (BioRad, Hercules, CA). All qPCR reactions were performed using the PowerUp™ SYBR Green Master Mix (Thermo Fisher Scientific, Waltham, MA). The qPCR reactions were performed using a StepOnePlus Real-Time PCR System (Applied Biosystems, Carlsbad, CA): UDG activation (50°C for 2 min), polymerase activation (95°C for 2 min), 40 cycles of denaturation (95°C for 15 sec) and annealing (various temperatures for 15 sec), extension (72°C for 60 sec). The primer sequences and annealing temperatures are provided in Table S3. Relative gene expression was determined using the Pfaffl method to account for differential primer efficiencies (Pfaffl, 2001), using the geometric mean of the Cq values for beta-2-microglobulin (*B2m*), GAPDH (*Gapdh*), and 18sRNA (*R18s*). The average Cq value from two livers from female C57/BL6J mice was used as the reference point. Data are reported as “Relative Expression.”

### Histological analyses

5μm liver sections were stained with hematoxylin and eosin. Micrographs (20x) were visualized on a Nikon Eclipse TE2000 microscope (Nikon Corporation; Tokyo, Japan).

### Statistical analyses

Data are presented as data points from individual mice or as means ± standard error (SE). Mice were considered hPPARα positive if they were either homozygous or heterozygous. Information on outliers is presented in Table S1. In the gene expression analyses, values more than four standard deviations different than the mean were excluded from the analyses. Overall, six values were excluded in five of the twenty genes analyzed. Within sex and genotype, statistical significance was determined by unpaired, two-tailed t-test (Prism 6, GraphPad Software Inc., La Jolla, CA). Regression analyses were performed using Microsoft R Open version 3.6.1. To investigate the interactions between treatment, sex and genotype in modifying phenotype and gene expression, we used multiple linear regression modeling (MLR). Each outcome was assessed using a MLR model with predictors including sex and an interaction term for genotype and treatment. Models were also stratified by sex, allowing effect estimates to vary between males and females. Statistical significance was evaluated at an α= 0.05 for all analyses.

## Results

Drinking water concentrations of PFOA averaged 3509 ± 138 μg/L. Based on average daily water consumption (0.21 ml/g mouse/day), the daily exposure was approximately 0.7 mg/kg/day. This resulted in serum concentrations of 47 ± 8 μg/ml in females and 48 ± 10 μg/ml in males (N = 4 for each sex). Assuming a 20 day half-life (Lou et al. 2009), and that at 6 weeks of exposure 77% of steady state was reached, the steady-state Cs/Cdw ≈ 18.

Daily exposure to PFOA for 6 weeks did not significantly impact weight gain in hPPARα mice of either sex (Fig. 1a and 1c). However, PFOA treatment significantly reduced weight gain in male PPARα null mice (Fig. 1c, Table 1). A similar trend was observed in female PPARα null mice, but the effect was not significant (Fig. 1a, Table 1). Body composition was not affected by PFOA in either genotype or sex (Fig. 1b and 1d). No differences in water or food consumption were observed (Fig. S1).

**Table 1:** Effect estimates (β) and standard errors (SE) for phenotypic outcomes. Regression models were fit to evaluate associations of phenotypic outcomes with treatment and genotype, including a treatment-genotype interaction term. The left hand column adjusts for sex. The two right columns stratify by sex, allowing results to differ between males and females. Statistical significance was evaluated at α = 0.05 for all analyses.

| Test | ALL |  | MALE |  | FEMALE |  |
| --- | --- | --- | --- | --- | --- | --- |
| | $\beta$ (SE) | P value | $\beta$ (SE) | P value | $\beta$ (SE) | P value |
| **Weight Gain, Week 6 (%)** |  |  |  |  |  |  |
| *PFOA treatment* | -24.62 (13.82) | 0.08 | -33.89 (22.10) | 0.14 | -15.43 (17.25) | 0.38 |
| *hPPARα Genotype* | -32.29 (12.65) | 0.01 | -47.58 (19.23) | 0.02 | -15.15 (16.63) | 0.37 |
| *Treatment\*Genotype* | 26.79 (18.11) | 0.15 | 43.54 (28.00) | 0.13 | 7.99 (23.51) | 0.74 |
| *Male Sex* | 25.71 (8.97) | 0.006 | - | - | - | - |
| **Body Fat (%)** |  |  |  |  |  |  |
| *PFOA treatment* | -2.87 (2.33) | 0.22 | -4.58 (3.76) | 0.23 | -1.28 (2.94) | 0.67 |
| *hPPARα Genotype* | -0.08 (2.13) | 0.97 | -1.64 (3.27) | 0.62 | 1.64 (2.83) | 0.57 |
| *Treatment\*Genotype* | 2.37 (3.05) | 0.44 | 4.96 (4.77) | 0.31 | -0.34 (4.00) | 0.93 |
| *Male Sex* | 0.89 (1.51) | 0.56 | - | - | - | - |
| **Liver to Body Weight (%)** |  |  |  |  |  |  |
| *PFOA treatment* | 4.75 (0.39) | <0.0001 | 4.99 (0.63) | <0.0001 | 4.49 (0.44) | <0.0001 |
| *hPPARα Genotype* | -0.41 (0.36) | 0.25 | -0.19 (0.54) | 0.72 | -0.60 (0.43) | 0.17 |
| *Treatment\*Genotype* | -1.17 (0.51) | 0.03 | -0.95 (0.79) | 0.24 | -1.50 (0.60) | 0.02 |
| *Male Sex* | 0.43 (0.25) | 0.10 | - | - | - | - |

**Fig. 1.**
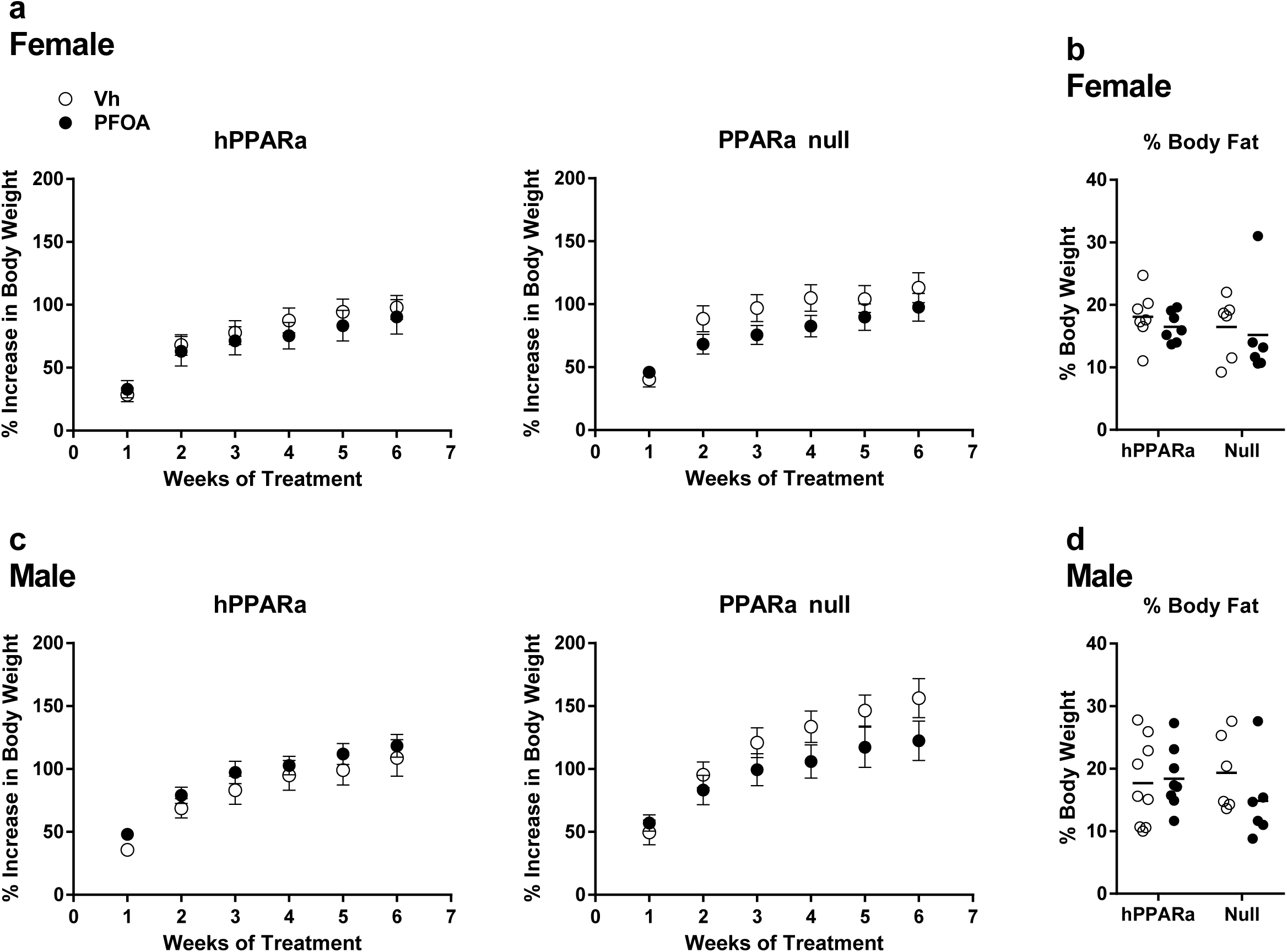
Weight gain in PFOA-exposed mice. Three-week-old male and female hPPARα and PPARα null mice were treated with either vehicle (Vh, NERL water with 5% sucrose) or PFOA (8 µM in NERL water with 5% sucrose) as drinking water for 6 weeks. During treatment, the mice were fed an American Diet (see Table S1). **a, c** Body weight (reported as percent increase from initial weight measured at weaning) was measured weekly. Data are presented as mean ± SE. N = 5-9. **b, d** Body composition was measured by EchoMRI. Data are from individual mice, with the mean indicated by a line. N = 5-9.

PFOA significantly increased liver to body weight ratios in both sexes and both genotypes (Fig. 2). In females, PFOA induced a significantly greater effect on liver to body weight ratios in PPARα null mice than in hPPARα mice (Fig. 2a, Table 1); this effect was not observed in males (Fig. 2c, Table 1). Histological analyses showed significant microvesicular lipid accumulation (steatosis) in PFOA-treated hPPARα mice of both sexes (Fig. 2b and 2d). Microvesicular steatosis also was evident in Vh-treated PPARα null mice of both sexes (Fig. 2b and 2d). Macrovesicular steatosis was present in PFOA-treated PPARα mice, with the largest lipid droplets observed in male PPARα null mice (Fig. 2b and 2d).

**Fig. 2.**
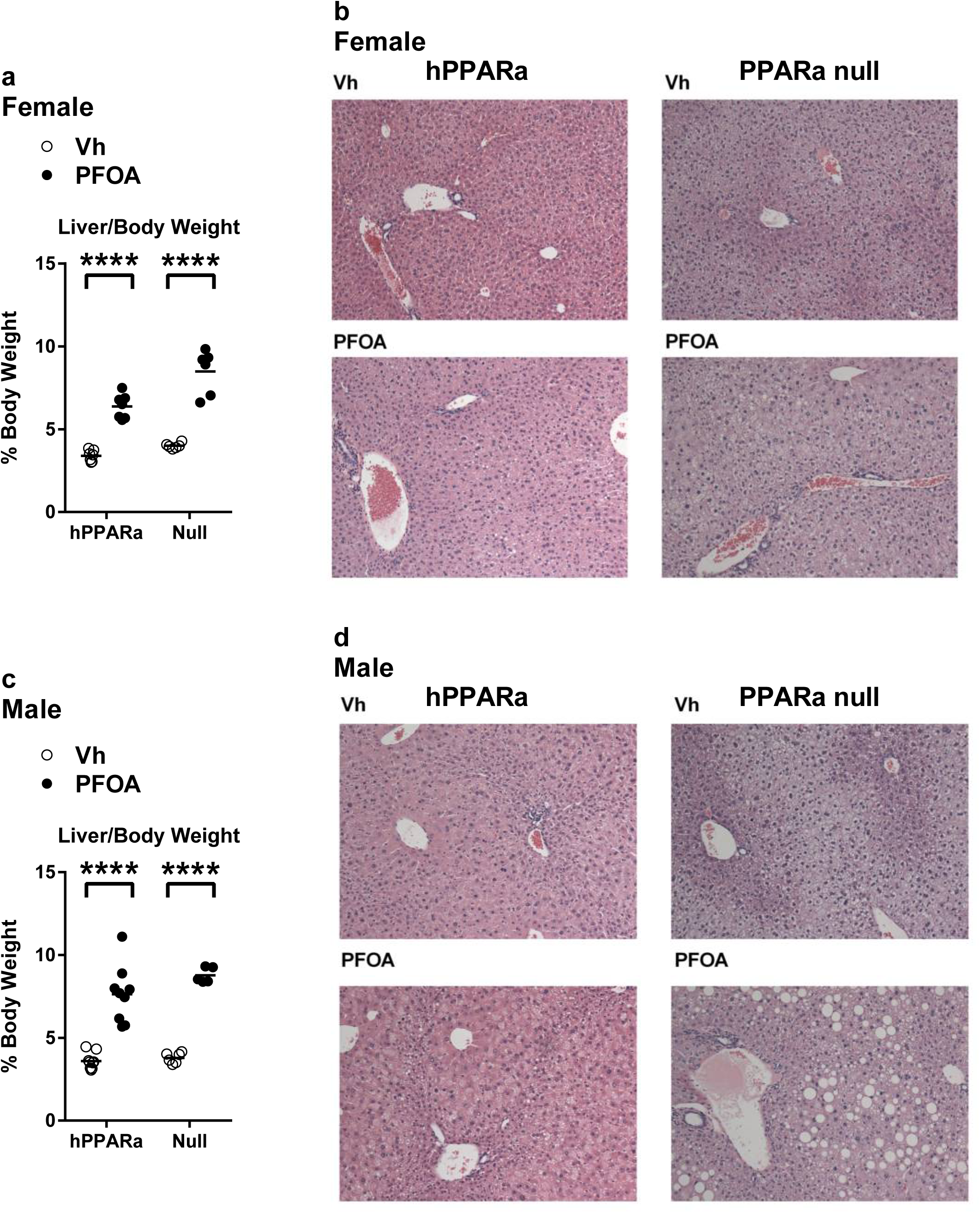
Liver/body weight in PFOA-exposed mice. hPPARα and PPARα null mice were exposed to Vh or PFOA in their drinking water for 6 weeks, as described in Fig. 1. **a, c** Liver to body weight ratios were determined at euthanasia. **b, d** H&E staining of representative liver sections. Data are from individual mice, with the mean indicated by a line. N = 5-9. Significantly different from Vh (**** p<0.0001, t-test).

Activation of hPPARα was evident in livers of PFOA-treated, humanized mice. Human *PPARA* mRNA was highly expressed in transgenic mice of both sexes, and lack of PPARα expression was confirmed in the PPARα null mice; expression of *PPARA* was not influenced by PFOA treatment (Fig. 3, Table 2). Expression of the PPARα target genes *Acox (*Acyl-CoA oxidase 1 is the first enzyme of the fatty acid beta-oxidation pathway), *Adrp* (Perlipin 2 coats intracellular lipid storage droplets), *Mogat1* (Monoacylglycerol O-acyltransferase 1 catalyzes the synthesis of diacylglycerols), and *Vnn1* (Vanin 1 biotransforms pantetheine in cysteamine and pantothenic acid, a precursor of coenzyme A) were upregulated by PFOA exposure in hPPARα mice but not in PPARα null mice in both sexes (Fig. 3, Table 2). *Pdk4* (Pyruvate dehydrogenase kinase 4 inhibits the pyruvate dehydrogenase complex) was upregulated by PFOA exposure in hPPARα mice but downregulated in female PPARα null mice (Fig. 3, Table 2). Sex-dependent differences in expression were evident for *Mogat1* and *Pdk4* (Table 2). *Mogat1* was upregulated to a greater extent by PFOA in male hPPARα mice. *Pdk4* was downregulated to a greater extent by PFOA in female hPPARα null mice.

**Table 2:**
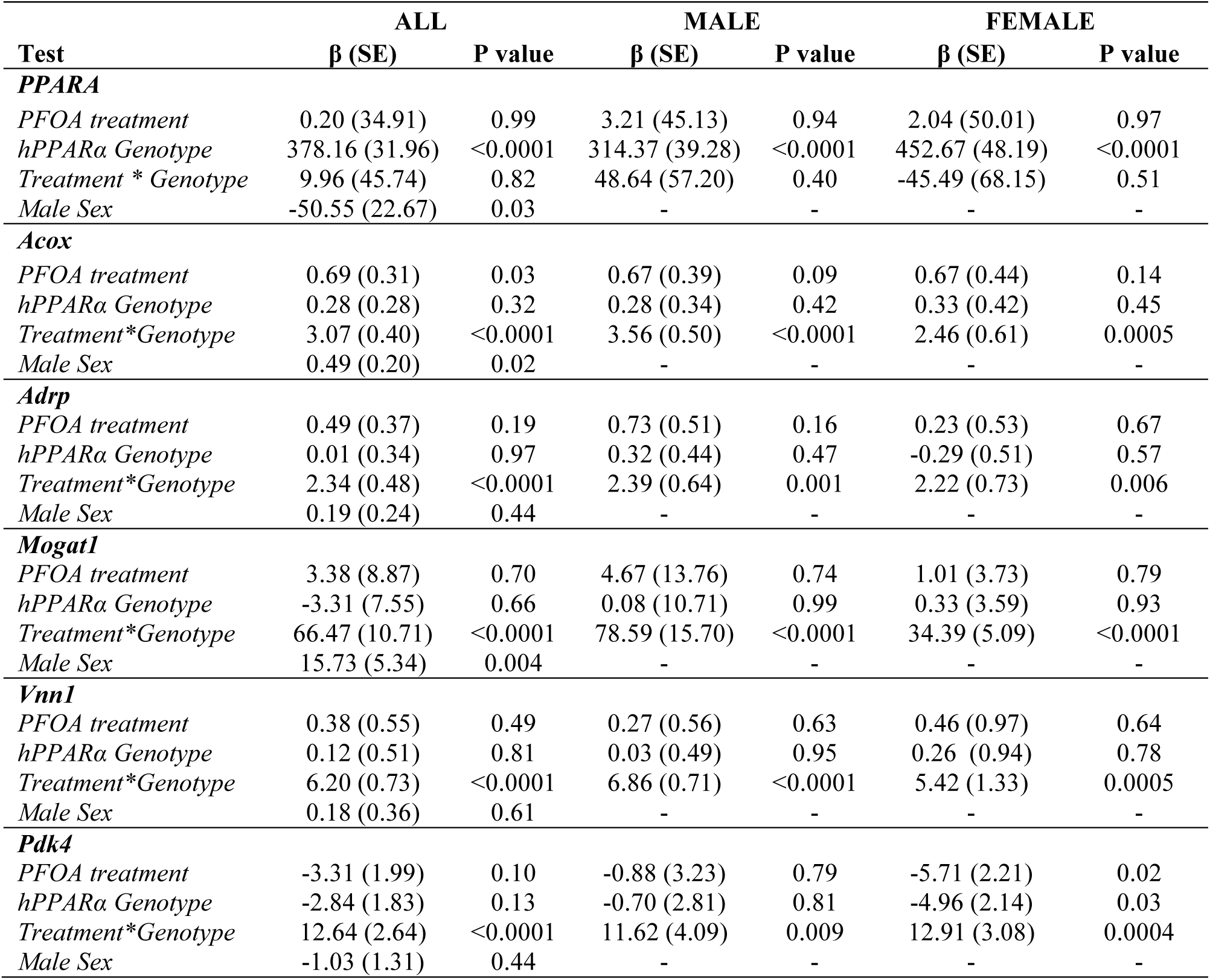
Effect estimates (β) and standard errors (SE) for relative expression of *PPARA* and its target genes. Regression models were fit to evaluate associations of gene expression outcomes with treatment and genotype, including a treatment-genotype interaction term. The left hand column adjusts for sex. The two right columns stratify by sex, allowing results to differ between males and females. Statistical significance was evaluated at α = 0.05 for all analyses.

**Fig. 3.**
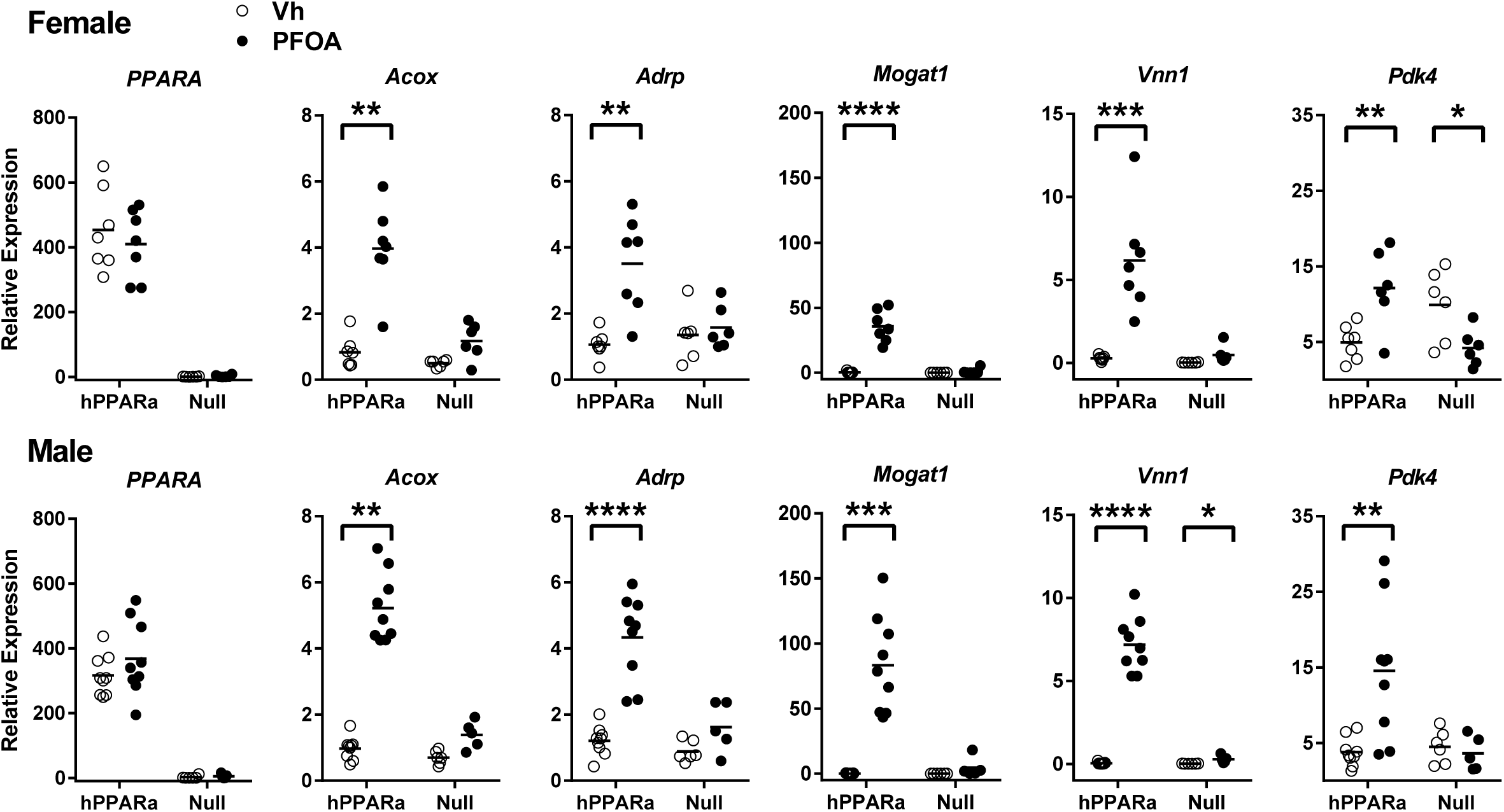
PPARα-target gene expression in liver of PFOA-exposed mice. hPPARα and PPARα null mice were exposed to Vh or PFOA in their drinking water for 6 weeks, as described in Fig. 1. Following isolation of RNA from liver, gene expression was determined by RT-qPCR. Data are from individual mice, with the mean indicated by a line. N = 5-9. Significantly different from Vh (* p<0.05, ** p<0.01, **** p<0.0001, t-test).

In addition to PPARα, evidence suggests that at least PPARγ and CAR also are molecular targets of PFAS (Abe et al. 2017; Buhrke et al. 2015; Vanden Heuvel et al. 2006). Expression of PPARγ mRNA (*Nr1c3*) along with its target genes *Fabp4* (Fatty acid binding protein 4 binds and transports long chain fatty acids) and *Cd36* (CD36 is a fatty acid translocase) were upregulated in PFOA treated mice of both sexes (Fig. 4a). There was a small but significant decrease in induction of *Nr1c3* expression and a trend toward a decrease in induction in *Fabp4* expression in male PPARα null mice (Table 3). Induction of *Cd36* was nearly completely abrogated in PPARα null mice of both sexes (Fig. 4a). In contrast, expression of CAR mRNA (*Nr1i3*) was modestly induced by PFOA in only male hPPARα mice (Fig. 4b). Expression of the CAR target genes *Cyp2b10* (Enzymes in the CYP2B family oxidatively metabolize a broad range endogenous and exogenous substrates) and *Gstm3* (GSTM3 is a glutathione s-transferase of the mu class) was highly upregulated in both sexes and in both genotypes (Fig. 4b). CAR target gene expression was induced to a greater extent in PPARα null mice than in hPPARα mice (Fig. 4b, Table 3).

**Table 3:** Effect estimates (β) and standard errors (SE) for relative expression of *Nr1c3* (PPARγ) and *Nr1i3* (CAR) and their target genes. Regression models were fit to evaluate associations of gene expression outcomes with treatment and genotype, including a treatment-genotype interaction term. The left hand column adjusts for sex. The two right columns stratify by sex, allowing results to differ between males and females. Statistical significance was evaluated at α = 0.05 for all analyses.

| Test | ALL |  | MALE |  | FEMALE |  |
| --- | --- | --- | --- | --- | --- | --- |
| | $\beta$ (SE) | P value | $\beta$ (SE) | P value | $\beta$ (SE) | P value |
| ***Nr1c3*** |  |  |  |  |  |  |
| *PFOA treatment* | 3.09 (0.64) | <0.0001 | 3.00 (0.98) | 0.005 | 3.16 (0.82) | 0.0009 |
| *hPPARα Genotype* | -0.89 (0.57) | 0.13 | -0.93 (0.85) | 0.29 | -0.79 (0.75) | 0.31 |
| *Treatment\*Genotype* | 2.55 (0.83) | 0.003 | 3.26 (1.24) | 0.01 | 1.67 (1.09) | 0.14 |
| *Male Sex* | -0.12 (0.41) | 0.77 | - | - | - | - |
| ***Fabp4*** |  |  |  |  |  |  |
| *PFOA treatment* | 0.39 (0.24) | 0.11 | 0.28 (0.31) | 0.38 | 0.48 (0.38) | 0.22 |
| *hPPARα Genotype* | 0.54 (0.22) | 0.02 | 0.57 (0.27) | 0.047 | 0.49 (0.36) | 0.19 |
| *Treatment\*Genotype* | 0.55 (0.31) | 0.09 | 0.68 (0.39) | 0.09 | 0.42 (0.52) | 0.42 |
| *Male Sex* | 0.08 (0.16) | 0.59 | - | - | - | - |
| ***Cd36*** |  |  |  |  |  |  |
| *PFOA treatment* | 1.50 (1.12) | 0.19 | 1.46 (1.62) | 0.38 | 1.55 (1.65) | 0.36 |
| *hPPARα Genotype* | 0.76 (1.00) | 0.45 | 0.64 (1.41) | 0.65 | 0.91 (1.51) | 0.55 |
| *Treatment\*Genotype* | 11.55 (1.45) | <0.0001 | 11.99 (2.05) | <0.0001 | 10.99 (2.19) | <0.0001 |
| *Male Sex* | -0.11 (0.72) | 0.88 | - | - | - | - |
| ***Nr1i3*** |  |  |  |  |  |  |
| *PFOA treatment* | -0.19 (0.14) | 0.20 | -0.28 (0.18) | 0.13 | -0.09 (0.21) | 0.66 |
| *hPPARα Genotype* | -0.07 (0.13) | 0.60 | -0.25 (0.16) | 0.12 | 0.15 (0.21) | 0.48 |
| *Treatment \* Genotype* | 0.29 (0.19) | 0.13 | 0.62 (0.23) | 0.01 | -0.11 (0.29) | 0.71 |
| *Male Sex* | -0.14 (0.09) | 0.15 | - | - | - | - |
| ***Cyp2b10*** |  |  |  |  |  |  |
| *PFOA treatment* | 54.00 (3.52) | <0.0001 | 43.51 (4.77) | <0.0001 | 63.09 (4.25) | <0.0001 |
| *hPPARα Genotype* | 0.35 (3.23) | 0.91 | 0.05 (4.15) | 0.99 | 0.16 (4.09) | 0.97 |
| *Treatment\*Genotype* | -27.13 (4.52) | <0.0001 | -16.83 (6.04) | 0.01 | -35.97 (5.79) | <0.001 |
| *Male Sex* | -4.09 (2.29) | 0.08 | - | - | - | - |
| ***Gstm3*** |  |  |  |  |  |  |
| *PFOA treatment* | 33.66 (3.67) | <0.0001 | 42.39 (5.94) | <0.0001 | 25.98 (3.74) | <0.0001 |
| *hPPARα Genotype* | 0.27 (3.35) | 0.93 | 0.86 (5.17) | 0.87 | 0.21 (3.60) | 0.95 |
| *Treatment\* Genotype* | -22.12 (4.79) | <0.0001 | -29.29 (7.53) | 0.0007 | -16.46 (5.09) | 0.004 |
| *Male Sex* | 4.84 (2.38) | 0.046 | - | - | - | - |

**Fig. 4.**
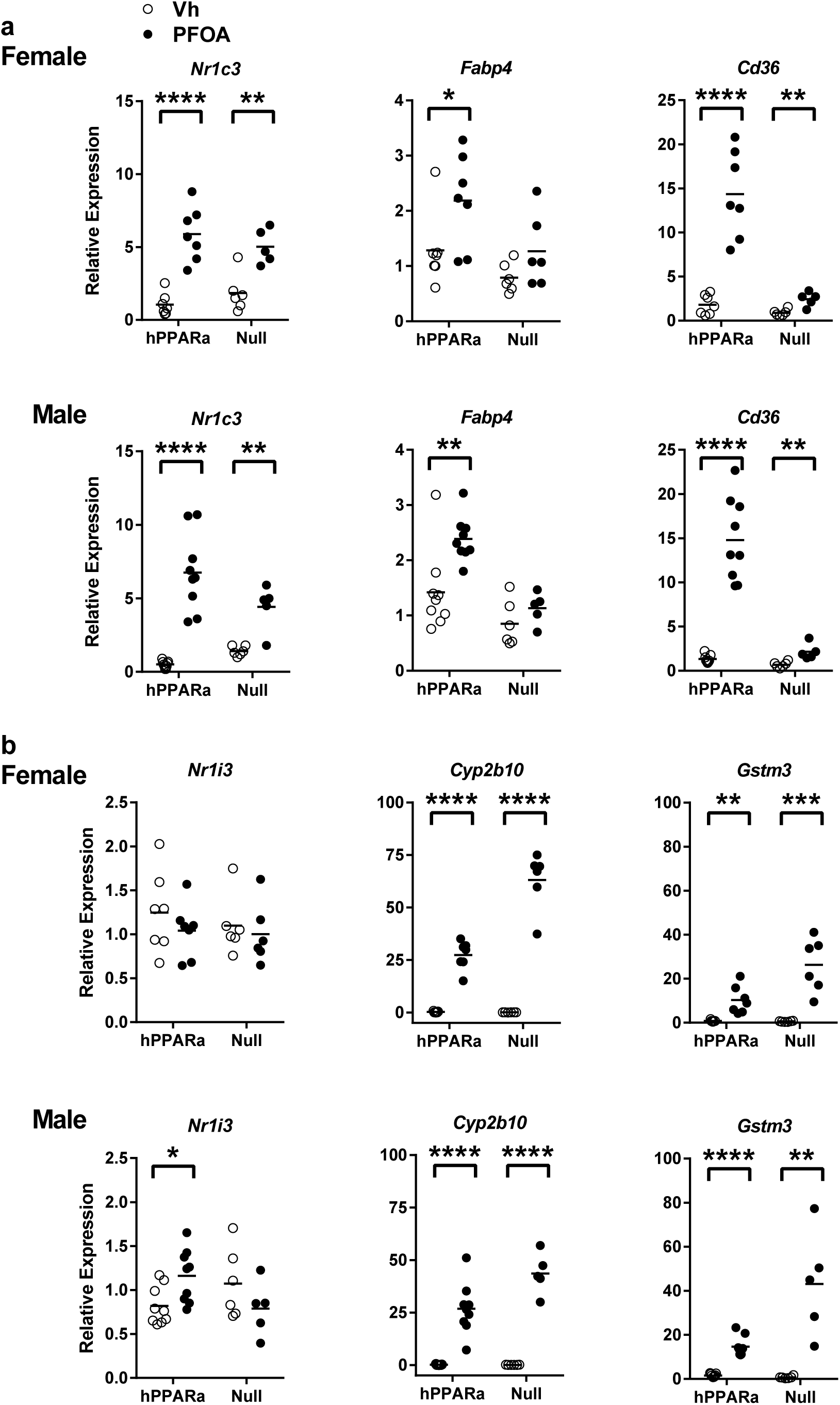
Alternative nuclear receptor-target gene expression in liver of PFOA-exposed mice. hPPARα and PPARα null mice were exposed to Vh or PFOA in their drinking water for 6 weeks, as described in Fig. 1. Following isolation of RNA from liver, gene expression was determined by RT-qPCR. **a** PPARγ (*Nr1c3*) and its target genes. **b** CAR (*Nr1i3*) and its target genes. Data are from individual mice, with the mean indicated by a line. N = 5-9. Significantly different from Vh (* p<0.05, ** p<0.01, *** p<0.001, **** p<0.0001, t-test).

Prior to experimentation, we hypothesized that activation of PPARα by PFOA may influence serum cholesterol homeostasis through four potential pathways in liver: increased *de novo* cholesterol synthesis, increased cholesterol export into the blood, decreased hepatic uptake of LDL-C from blood, and/or decreased cholesterol turnover to bile acids (Fig. 5). Expression of *Hmgcr*, the rate limiting step in cholesterol synthesis, was decreased by PFOA exposure in hPPARα but not PPARα null mice (Fig. 5a, Table 4). Expression of *Apob*, the apolipoprotein associated with VLDL-C and LDL-C, was not changed by PFOA exposure in either genotype or sex (Fig. 5b). Expression of *Ldlr*, which is responsible for hepatic uptake of LDL-C, was decreased by PFOA exposure in both hPPARα and PPARα null mice (Fig. 5c). Lastly, expression of *Cyp7a1*, the rate limiting step in conversion of cholesterol to bile acids and thus cholesterol efflux from the body, was down regulated by PFOA in both hPPARα and PPARα null mice but more so in hPPARα mice than PPARα null mice (Fig. 5d). Interestingly, PFOA’s effect on expression of cholesterol homeostasis genes, particularly *Cyp7a1*, was greater in female than male mice (Table 4).

**Table 4:** Effect estimates (β) and standard errors (SE) for relative expression of genes contributing cholesterol homeostasis. Regression models were fit to evaluate associations of gene expression outcomes with treatment and genotype, including a treatment-genotype interaction term. The left hand column adjusts for sex. The two right columns stratify by sex, allowing results to differ between males and females. Statistical significance was evaluated at α = 0.05 for all analyses.

| Test | ALL |  | MALE |  | FEMALE |  |
| --- | --- | --- | --- | --- | --- | --- |
| | $\beta$ (SE) | P value | $\beta$ (SE) | P value | $\beta$ (SE) | P value |
| ***Hmgcr*** |  |  |  |  |  |  |
| *PFOA treatment* | 0.04 (0.10) | 0.68 | 0.11 (0.16) | 0.50 | -0.01 (0.14) | 0.93 |
| *hPPARα Genotype* | 0.21 (0.10) | 0.04 | 0.17 (0.14) | 0.21 | 0.26 (0.14) | 0.08 |
| *Treatment\*Genotype* | -0.32 (0.14) | 0.03 | -0.29 (0.20) | 0.16 | -0.38 (0.20) | 0.07 |
| *Male Sex* | -0.03 (0.07) | 0.61 | - | - | - | - |
| ***ApoB*** |  |  |  |  |  |  |
| *PFOA treatment* | 0.20 (0.16) | 0.19 | -0.04 (0.20) | 0.84 | 0.42 (0.24) | 0.09 |
| *hPPARα Genotype* | 0.13 (0.15) | 0.40 | 0.14 (0.18) | 0.44 | 0.08 (0.24) | 0.73 |
| *Treatment\*Genotype* | -0.08 (0.21) | 0.70 | 0.07 (0.26) | 0.78 | -0.18 (0.33) | 0.60 |
| *Male Sex* | 0.10 (0.10) | 0.34 | - | - | - | - |
| ***Ldlr*** |  |  |  |  |  |  |
| *PFOA treatment* | -0.11 (0.05) | 0.03 | -0.03 (0.08) | 0.73 | -0.19 (0.05) | 0.0004 |
| *hPPARα Genotype* | -0.04 (0.04) | 0.31 | 0.005 (0.07) | 0.95 | -0.09 (0.04) | 0.047 |
| *Treatment\*Genotype* | 0.03 (0.06) | 0.64 | -0.005 (0.10) | 0.96 | 0.05 (0.06) | 0.46 |
| *Male Sex* | -0.02 (0.03) | 0.49 | - | - | - | - |
| ***Cyp7a1*** |  |  |  |  |  |  |
| *PFOA treatment* | -0.63 (0.20) | 0.003 | -0.55 (0.31) | 0.09 | -0.68 (0.21) | 0.004 |
| *hPPARα Genotype* | 0.69 (0.19) | 0.0005 | 0.34 (0.28) | 0.23 | 1.08 (0.20) | <0.0001 |
| *Treatment\*Genotype* | -0.31 (0.27) | 0.26 | 0.02 (0.40) | 0.96 | -0.76 (0.29) | 0.02 |
| *Male Sex* | -0.30 (0.13) | 0.03 | - | - | - | - |

**Fig. 5.**
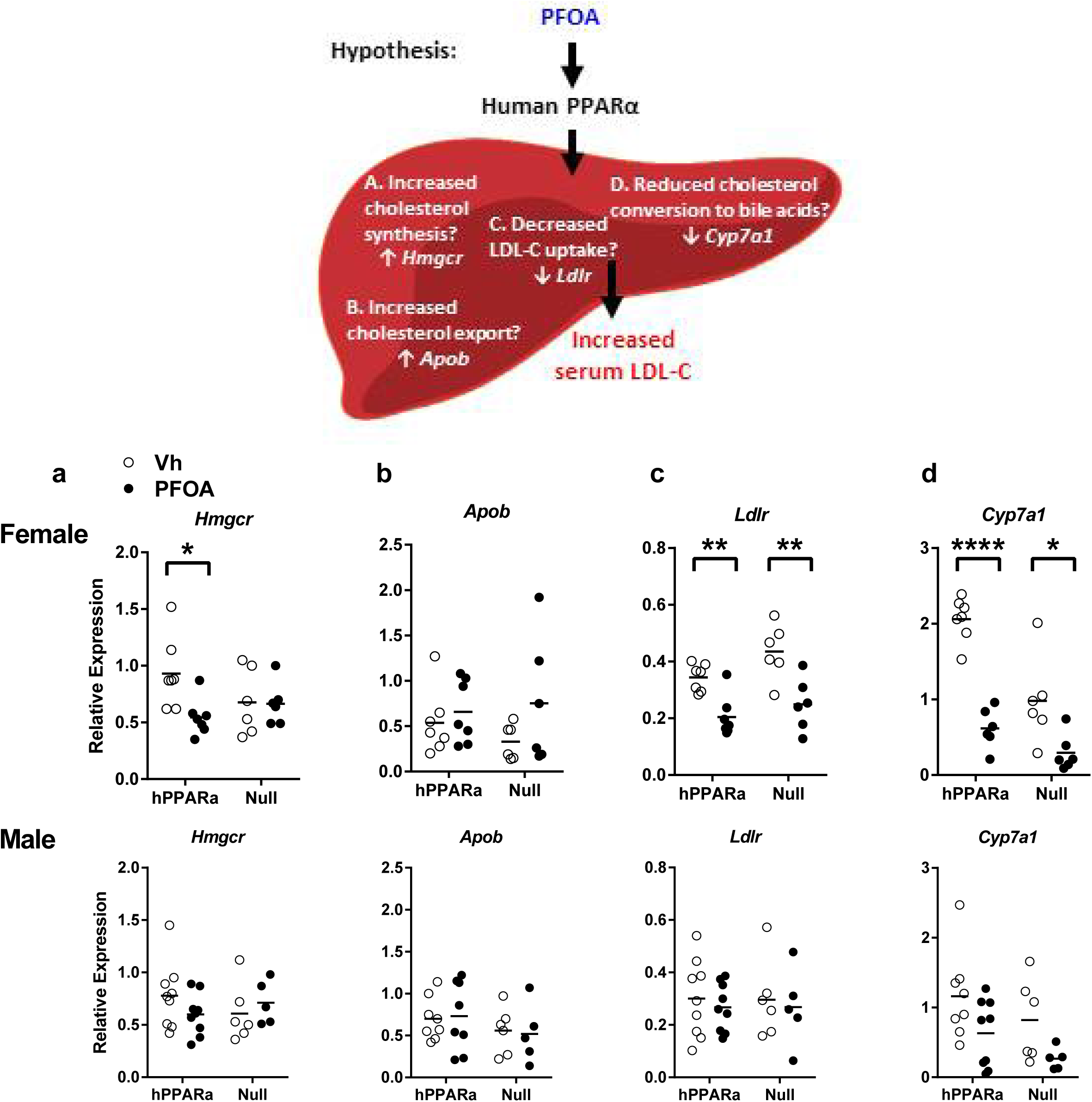
Cholesterol homeostasis-related gene expression in liver of PFOA-exposed mice. hPPARα and PPARα null mice were exposed to Vh or PFOA in their drinking water for 6 weeks, as described in Fig. 1. Following isolation of RNA from liver, gene expression was determined by RT-qPCR. The hypothetical model indicates biomarker genes for each of the pathways. **a** Cholesterol synthesis. **b** Cholesterol export. **c** Cholesterol import. **d** Cholesterol efflux. Data are from individual mice, with the mean indicated by a line. N = 5-9. Significantly different from Vh (* p<0.05, ** p<0.01, *** p<0.001, **** p<0.0001, t-test).

While PPARα is known to regulate cholesterol homeostasis (Bouchard-Mercier et al. 2011; Flavell et al. 2000; Robitaille et al. 2004; Sparso et al. 2007; Tanaka et al. 2007; Vohl et al. 2000), there are no studies showing direct interactions of PPARα with the promoters of the genes involved in cholesterol homeostasis. Rather, *Hmgcr* is regulated by SREBP2 (*Srebf2*; (Brown and Goldstein 1997; Sharpe and Brown 2013)). *Apob* is regulated by C/EBPα and HNF4α (Metzger et al. 1993). *Ldlr* is regulated by SREBP1 (*Srebf1*), SREBP2 and estrogen receptor (Brown and Goldstein 1997; Parini et al. 1997; Yokoyama et al. 1993). *Cyp7a1* is regulated by HNF4α, LXR and FXR (Gupta et al. 2002; Kir et al. 2012). PFOA treatment did not regulate C/EBPα, HNF4α, SREBP1 or SREBP2 at the transcriptional level (Fig. 6), although regulation at the post-translational level is still a possibility. Interestingly, *Srebf1* was significantly downregulated in PPARα null mice compared to hPPARα mice (Fig. 6, Table S4)

**Fig. 6.**
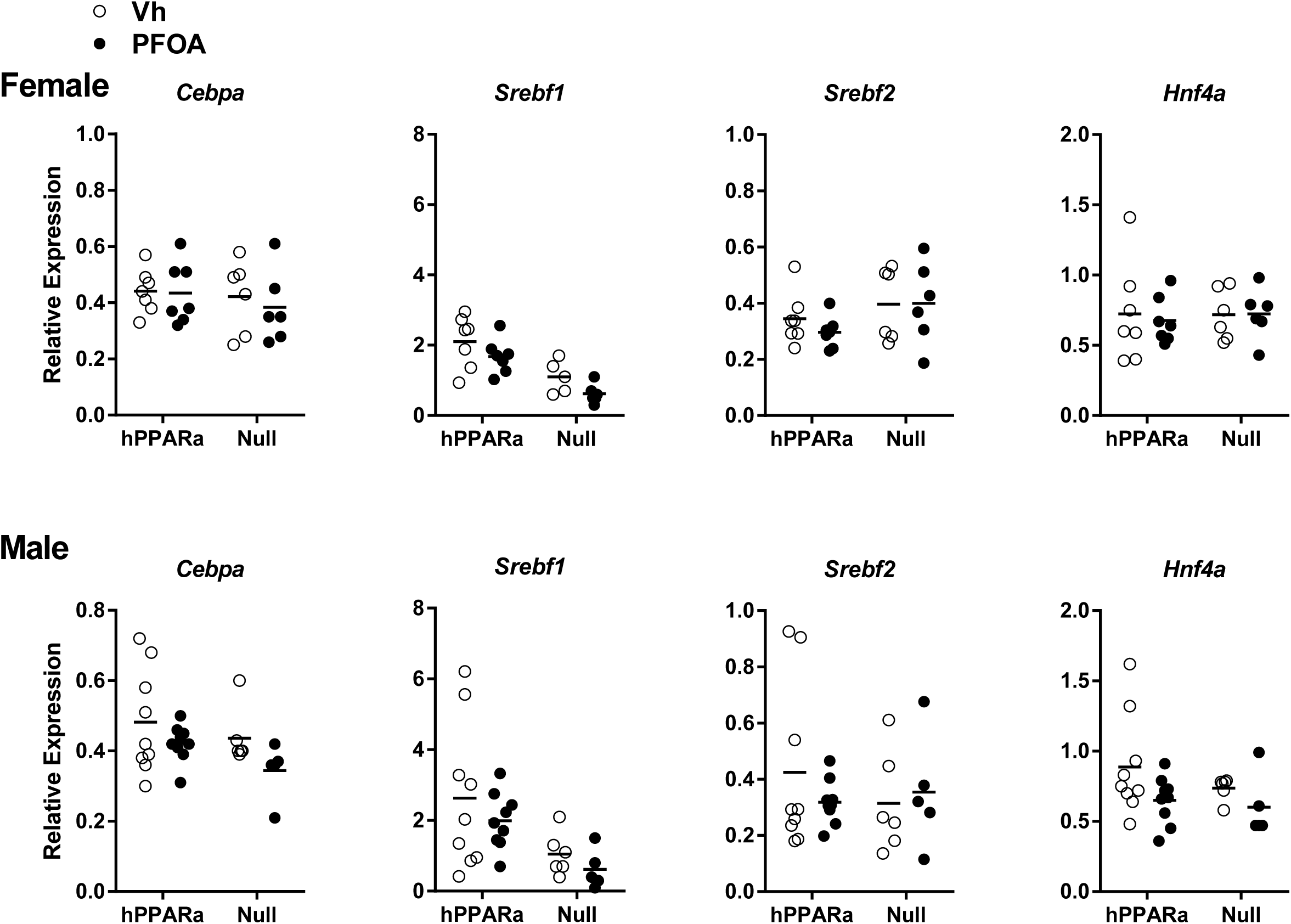
Expression of transcription factors that regulate cholesterol homeostasis in liver of PFOA-exposed mice. hPPARα and PPARα null mice were exposed to Vh or PFOA in their drinking water for 6 weeks, as described in Fig. 1. Following isolation of RNA from liver, gene expression was determined by RT-qPCR. Data are from individual mice, with the mean indicated by a line. N = 5-9. No significant differences were detected (t-test).

## Discussion

Increased serum total cholesterol and LDL-C are strongly associated with PFAS exposure in humans. However, studies of the effects of PFOA on serum lipids and cholesterol in animal models have produced contradictory results. New models are needed to investigate the mechanism(s) of action through which PFAS could interfere with cholesterol homeostasis. Here, we tested the hypothesis that PPARα activation is a critical molecular initiating event in the adverse outcome pathway linking human-relevant PFOA exposure with dyslipidemia in a novel model, mice expressing hPPARα fed an American diet. Over all, PFOA modulates at least the PPARα, PPARγ and CAR pathways in hPPARα mice, as well as multiple genes involved in cholesterol metabolism and homeostasis. Not all effects were PPARα-dependent. Our results show that the hepatic response to PFOA exposure is sexually dimorphic.

The PFOA exposure in this study was designed to recapitulate serum PFOA concentrations observed in some epidemiological studies. The PFOA’s toxicokinetic parameters differ significantly between rodents and humans with rodents having increased renal clearance capacity and a substantially shorter half-life compared with humans (Harada et al. 2007; Lou et al. 2009). Thus, we used a higher than typical exposure dose (≈ 0.7 mg/kg/day) to generate a serum PFOA concentration (≈ 48 μg/ml) similar to that found in fluorochemical workers in the US (≈ 22 μg/ml (Steenland et al. 2010)). We estimated a steady-state serum (Cs) to drinking water (Cdw) ratio of ≈ 18 in the mice in this study, which is similar to the ratio previously determined in CD1 mice (Cs/Cdw ≈ 12 (White et al. 2011)). As expected from the differences in half life, in humans the Cs/Cdw ratio was estimated to be on the order of 114 and 141 (Hoffman et al. 2011).

Our results demonstrate that both hPPARα and PPARα null mice exposed to PFOA and fed an American diet develop hepatosteatosis. PFOA induced a significant increase in liver:body weight ratio associated with histologically evident increases in hepatocyte lipid accumulation. PFOA-induced hepatosteatosis has been observed previously in mice expressing wildtype PPARα (mPPARα), hPPARα mice and PPARα null mice (Das et al. 2017; Minata et al. 2010; Nakagawa et al. 2012; Nakamura et al. 2009; Tan et al. 2013). While hepatosteatosis is induced in PFOA-exposed mice fed standard composition rodent diets, the severity is increased when mice are co-exposed to PFOA and a high fat diet (Tan et al., 2013). Importantly, in an exposure scenario that generated an approximately steady state body burden, hPPARα mice were more susceptible to hepatic steatosis than mPPARα mice (Nakagawa et al. 2012), underscoring the importance of using a humanized mouse model to investigate PFOA-induced hepatic endpoints.

PFOA activated hPPARα. However, upregulation of PPARγ and CAR target genes indicate that PFOA exerts biological effects through multiple pathways. The ability of PFOA to upregulate the transcription of PPARγ has been documented (Rosen et al. 2008). Upregulation of *Fabp4* and *Cd36*, genes classically thought to be regulated by PPARγ was abrogated in PPARα null mice. This is not necessarily a surprise, as there can be a significant overlap in regulation of genes by PPARs. While regulation of fatty acid transporter genes is largely controlled by PPARγ in adipose, their expression is PPARα-dependent in liver (Motojima et al. 1998). CAR transcriptional activity was strongly upregulated by PFOA in both hPPARα and PPARα null mice. Studies with PXR, CAR and FXR null mice show that CAR is the most significant contributor to PFAS-induced changes in gene expression, after PPARα (Abe et al. 2017; Cheng and Klaassen 2008). Interestingly, our findings show that PFOA induces CAR target gene expression to a higher degree in PPARα null mice than hPPARα mice indicating a potential interaction between the two nuclear receptors. Increased CAR activation in the absence of PPARα has been identified previously and may be due to antagonistic effects of these two nuclear receptors (Corton et al. 2014; Rosen et al. 2017).

We hypothesized that PFOA alters lipid homeostasis through one of four mechanisms in the liver: increased *de novo* cholesterol synthesis, increased cholesterol export into the blood, decreased hepatic uptake of LDL-C from blood, and/or decreased cholesterol turnover to bile acids. We observed a significant reduction in the expression of *Hmgcr, Ldlr*, and *Cyp7a1* in female mice. *Apob* expression was unchanged in both sexes. These changes in expression also were observed in C57BL/6J female mice (Rebholz et al. 2016). While the decrease in *Hmgcr* expression would be expected to decrease serum cholesterol, reductions in *Cyp7a1* and *Ldlr* expression would be expected in increase serum cholesterol. The downregulation of *Cyp7a1* occurs more broadly across models, as we observed *Cyp7a1* downregulation in both male and female SV129 mice and Rebholz et al., 2016 observed its downregulation in both sexes in both C57Bl/6J and Balb/C mice. Importantly, PFOA also downregulates *Cyp7a1* expression in human hepatocytes (Behr et al. 2020). Cholesterol is converted to bile acids, a major mechanism for removal of cholesterol from the body, through two primary molecular pathways with CYP7A1 being the rate limiting enzyme in the primary pathway (Dietschy and Turley 2002). In humans, the CYP7A1-mediated primary pathway accounts for 90% of bile acid production whereas it only accounts for 60% in mice (Phelps et al. 2019). Further, CYP7A1 is a major factor regulating lipoprotein synthesis and assembly (Wang et al. 1997). A reduction of more than 50% in *Cyp7a1* expression was associated with increased plasma cholesterol levels in mice (Rebholz et al., 2016). Expression of *Ldlr* plays a substantial role in regulating serum LDL-C, with serum concentrations increasing an order of magnitude in *Ldlr* null mice (Osono et al. 1995). It is likely then that the PFOA-induced decrease in *Ldlr* expression has a biologically significant effect on serum cholesterol. Intriguingly, PFOA substantially upregulated *Vnn1*, overexpression of which is associated with increased liver lipid content, serum triglycerides and LDL-C, decreased HDL-C and enhanced atherosclerotic plaque formation (Hu et al. 2016). It remains to be determined if any or all of these changes in gene expression contribute to serum lipid dysregulation of PFOA.

Only the repression of *Hmgcr* expression by PFOA appeared to be PPARα dependent. *Hmgcr* expression was also decreased in PPARα null mice. Previous studies have shown that GW7647, a PPARα ligand, increases the occupancy of PPARα at the *Hmgcr* promoter, in concert with SREBPs (van der Meer et al. 2010); however this resulted in an increased expression of *Hmgcr*. Thus PFOA-liganded PPARα appears to be acting distinctly from GW7647-liganded PPARα. On the other hand, repression of *Ldlr* and *Cyp7a1* expression by PFOA occurred in both hPPARα and PPARα null mice. PPARα overexpression or treatment with a PPARα ligand has been shown to suppress HNF4α protein expression, thereby reducing its interaction with the *Cyp7a1* promoter (Marrapodi and Chiang 2000). However, we did not observe a decrease in *Hnf4a* mRNA in PFOA-treated animals. No studies have reported regulation of *Ldlr* by PPARα.

A single previous study has investigated the effects of PFAS on lipid homeostasis in female mice (Rebholz et al. 2016). The results presented here corroborate the observation of sex-dependent effects of PFOA on liver physiology. The data show differences at the macro level (liver:body weight) and gene expression level. Most interesting are the differences in effect of PFOA on expression of genes involved in cholesterol homeostasis. It is well known that cholesterol homeostasis differs in mouse strains and sexes (Bruell et al. 1962). However, before the current study, only Rebholz et al., 2016 investigated the influence of strain and sex on the response to PFOA and showed that C57Bl/6J mice were more sensitive to modulation of cholesterol homeostasis by PFOA than Balb/C mice. They also showed that female C57BL/6 mice, the mice with the greatest increase in serum cholesterol, were the only mice to have significant changes in expression of multiple genes involved in cholesterol homeostasis (Rebholz et al. 2016). We observed significant changes due to PFOA in these same genes (*Hmgcr, Ldlr*, and *Cyp7a1*) in female hPPARα mice. It is critical that future studies take into account the complexity of the genetics that contribute to cholesterol homeostasis when investigating PFOA-induced effects.

## Conclusions

PFOA activates human PPARα and CAR at human relevant serum concentrations *in vivo*. Multiple genes involved in cholesterol homeostasis are modified by PFOA by both PPARα-dependent and independent mechanisms. Investigation of the effects of PFOA and their dependence on PPARα beyond the four biomarker genes analyzed here is necessary. The essential role of hPPARα in basal cholesterol homeostasis, as well as fatty acid homeostasis, and known species differences in ligand binding gene batteries support the conclusion that our model is an important new tool in dissecting the multiple, interacting mechanisms of PFOA action on cholesterol homeostasis. Importantly, PFOA-induced effects appear to be stronger in females than in males. Regulation of cholesterol homeostasis is complex, is modified by diet, with multiple pathways able to compensate for deficiencies (Dietschy and Turley 2002; Dietschy et al. 1993). Thus, further research is needed to delineate the biologically significant effects of PFAS on multiple aspects of cholesterol homeostasis.

## Acknowledgements

This work was supported by the National Institute of Environmental Health Sciences Superfund Research Program P42 ES007381 (Jennifer Schlezinger) and R01 ES027813 (Thomas Webster). Greylin Nielsen and Jennifer Oliver are supported by training grant T32 ES01456. The authors thank Mr. Nathan Burritt for his excellent technical assistance and Dr. Juliet Gentile (Research Diets, Inc.) for her expert assistance in designing the diet.

