## Supplemental Material for "Perfluorooctanoic acid activates multiple nuclear receptor pathways and skews expression of genes regulating cholesterol homeostasis in liver of humanized PPARα mice fed an American diet"

**Table of Contents**

| | | Page |
|---|---|---|
| Table S1 | Cohort information. | 2 |
| Table S2 | Components of the "What we eat in America" diet (Research Diets D18110214) | 3 |
| Table S3 | Mouse and human primer sequences for reverse transcriptase qPCR. | 4 |
| Table S4 | Effect estimates (β) and standard errors (SE) for expression of transcription factors regulating cholesterol homeostasis. | 5 |
| Fig. S1 | Drinking water (a) and food (b) consumption in PFOA-treated mice. | 6 |

**Table S1: Cohort information**

| | | *PPARA* | | | | Knockout | | | |
|---|---|---|---|---|---|---|---|---|---|
| | | Vh | | PFOA | | Vh | | PFOA | |
| **Cohort Number** | **Breeders** | **Male Pups** | **Female Pups** | **Male Pups** | **Female Pups** | **Male Pups** | **Female Pups** | **Male Pups** | **Female Pups** |
| 1 | Male 1/ Female 1 | 1 | 2 | | | | | | |
| 2 | Male 2/ Female 2 | 1 | | 1 | 1 | 1 | 1 | | 1 |
| 3 [a] | Male 1/ Female 1 | 1 | 1 | | | 1 | 1 | | |
| 4 | Male 2/ Female 2 | | | 1 | | | | 1 | 1 |
| 5 | Male 1/ Female 1 | | | 1 | 2 | | | 2 | 2 |
| 6 | Male 2/ Female 2 | 2 | 2 | 1 | | 1 | 3 | | |
| 7 | Male 1/ Female 1 | 3 | | | 1 | 1 | | | |
| 8 [b] | Male 2/ Female 2 | | 1 | 1 | 2 | | | | 1 |
| 9 | Male 2/ Female 1 | | | | | 2 | 1 | 2 | 1 |
| 10 | Male 4/ Female 4 | | 1 | 2 | | | | | |
| 11 [c] | Male 2/ Female 3 | 1 | | 1 | 1 | | | 1 | |
| | **Total** | **9** | **7** | **9** | **7** | **6** | **6** | **5** | **6** |

[a]One female failed to thrive (the final body weight was more than 2 standard deviations lower than the average final body weight) and was excluded from data analysis.
[b]One female had total serum lipids 4 standard deviations above the mean and was excluded from data analysis.
[c]One male had a severe eye injury and was euthanized early.

**Table S2: Components of the "What we eat in America" diet (Research Diets D18110214).**

| Macronutrient | % Grams | % kcal |
|---|---|---|
| Protein | 16 | 14.7 |
| Carbohydrate | 57 | 51.8 |
| Fat | 16 | 33.5 |
| Fiber | 5.4 | |
| Cholesterol | 0.05 | |
| Kcal/gm | 4.4 | |
| **Ingredient** | **grams** | **kcal** |
| Casein | 146 | 584 |
| L-cysteine | 3 | 12 |
| Corn Starch | 176.81 | 707 |
| Maltodextrin 10 | 100 | 400 |
| Sucrose | 238.67 | 955 |
| Cellulose | 40 | |
| Inulin | 10 | |
| Soybean Oil | 25 | 225 |
| Lard | 75.6 | 680 |
| Butter | 50.4 | 454 |
| Mineral Mix S10026 | 10 | |
| Dicalcium Phosphate | 13 | |
| Calcium Carbonate | 5.5 | |
| Potassium Citrate Monohydrate | 16.5 | |
| Vitamin Mix V10001 | 10 | 40 |
| Choline Bitartrate | 2 | |
| Cholesterol | 0.2943 | |
| FD&C Yellow Dye #5 | 0.025 | |
| FD&C Red Dye #40 | 0.025 | |
| **Total** | 922.8243 | 4057 |

**Table S3. Mouse (M) and human (H) primer sequences for reverse transcriptase qPCR.**

| Gene Symbol | FORWARD | REVERSE | Annealing Temp. °C |
|---|---|---|---|
| *M-R18s* | GTAACCCGTTGAACCCCATT | CCATCCAATCGGTAGTAGCG | 55 |
| *M-B2m* | CTGCTACGTAACACAGTTCCACCC | CATGATGCTTGATCACATGTCTCG | 55 |
| *M-Gapdh* | ACAGTCCATGCCATCACTGCC | GCCTGCTTCACCACCTTCTTG | 55 |
| *M-Pdk4* | TTTCTCGTCTCTACGCCAAG | GATACACCAGTCATCAGCTTCG | 58 |
| *M-Acox* | AGCGAGCCAGAGCCCCAG | TCAGGCAGCTCACTCAGG | 59 |
| *M-Gstm3* | TATGACACTGGGCTATTGGAACA | GGGCATCCCCCATGAC | 58 |
| *M-Srebf1* | GCGTTCTGGAGACCATGGA | ACAAAGTTGCTCTGAAAACAAATCA | 58 |
| *M-Hnf4a* | AGAGGTTCTGTCCCAGCAGATC | CGTCTGTGATGTTGGCAATC | 57 |
| *M-Hmgcr* | AGCTTGCCCGAATTGTATGTG | TCTGTTGTGAACCATGTGACTTC | 60 |
| *M-Apob* | CAGGTGGCCACAGCCAATAA | ACTGCAGGTCTGGCTCAGGA | 61 |
| *M-Ldlr* | CTAGCAAGTGGGTGTGCGATG | CTGAATTGATTGGACTGACAGGTGA | 59 |
| *M-Cyp7a1* | TGCCTTCTGCTACCGAGTGA | GGGCTTTATGTGCGGTCTTG | 59 |
| *M-Adrp* | TCCACTGTCCACCTGATTGA | TGGCATGTAGTCTGGAGCTG | 60 |
| *M-Nr1c3 (Pparg1/2)* | GACCACTCGCATTCCTTT | CCACAGACTCGGCACTCA | 55 |
| *M-Fabp4* | GATGAAATCACCGCAGACG | GCCCTTTCATAAACTCTTGTGG | 55 |
| *M-Cd36* | GGTGATGTTTGTTGCTTTTATGATTTC | TGTAGATCGGCTTTACCAAAGATG | 54 |
| *M-Cyp2b10* | AAAGTCCCGTGGCAACTTCC | TCCCAGGTGCACTGTGAACA | 61 |
| *M-Nr1i3 (Car)* | CTCAAGGAAAGCAGGGTCAG | AGTTCCTCGGCCCATATTCT | 58 |
| *M-Mogat1* | TGGTGCCAGTTTGGTTCCAG | TGCTCTGAGGTCGGGTTCA | 61 |
| *M-Vnn1* | TATGTCTTCCCTGAAGTGTT | CCCAGTCCTTCCCATAC | 54 |
| *H-PPARA* | CAGAACAAGGAGGCGGAGGTC | TTCAGGTCCAAGTTTGCGAAGC | 55 |

**Table 4: Effect estimates (β) and standard errors (SE) for expression of transcriptional factors that regulate cholesterol homeostasis**

Regression models were fit to evaluate associations of gene expression outcomes with treatment and genotype, including a treatment-genotype interaction term. The left hand column adjusts for sex. The two right columns stratify by sex, allowing results to differ between males and females. Statistical significance was evaluated at α = 0.05 for all analyses.

| | ALL | | MALE | | FEMALE | |
|---|---|---|---|---|---|---|
| **Test** | **β (SE)** | **P value** | **β (SE)** | **P value** | **β (SE)** | **P value** |
| ***Cebpa*** | | | | | | |
| *PFOA treatment* | -0.04 (0.04) | 0.31 | -0.09 (0.06) | 0.14 | -0.003 (0.07) | 0.96 |
| *hPPARα Genotype* | 0.03 (0.04) | 0.40 | 0.04 (0.05) | 0.41 | 0.02 (0.06) | 0.77 |
| *Treatment*Genotype* | 0.01 (0.06) | 0.91 | 0.03 (0.08) | 0.69 | -0.004 (0.09) | 0.96 |
| *Male Sex* | -0.002 (0.03) | 0.92 | - | - | - | - |
| ***Srebf1*** | | | | | | |
| *PFOA treatment* | -0.42 (0.42) | 0.33 | -0.39 (0.79) | 0.62 | -0.49 (0.32) | 0.14 |
| *hPPARα Genotype* | 1.33 (0.39) | 0.001 | 1.60 (0.69) | 0.03 | 1.00 (0.31) | 0.004 |
| *Treatment*Genotype* | -0.13 (0.25) | 0.82 | -0.24 (1.00) | 0.81 | 0.06 (0.43) | 0.88 |
| *Male Sex* | 0.24 (0.28) | 0.39 | - | - | - | - |
| ***Srebf2*** | | | | | | |
| *PFOA treatment* | 0.02 (0.07) | 0.74 | 0.04 (0.13) | 0.75 | 0.002 (0.06) | 0.97 |
| *hPPARα Genotype* | 0.03 (0.06) | 0.60 | 0.11 (0.11) | 0.33 | -0.05 (0.06) | 0.40 |
| *Treatment*Genotype* | -0.10 (0.09) | 0.26 | -0.15 (0.16) | 0.37 | -0.05 (0.09) | 0.56 |
| *Male Sex* | 0.002 (0.05) | 0.96 | - | - | - | - |
| ***Hnf4a*** | | | | | | |
| *PFOA treatment* | -0.06 (0.10) | 0.56 | -0.13 (0.15) | 0.37 | 0.003 (0.14) | 0.98 |
| *hPPARα Genotype* | 0.09 (0.09) | 0.35 | 0.15 (0.13) | 0.25 | 0.01 (0.13) | 0.96 |
| *Treatment*Genotype* | -0.09 (0.13) | 0.48 | -0.10 (0.19) | 0.59 | -0.05 (0.19) | 0.80 |
| *Male Sex* | 0.02 (0.07) | 0.76 | - | - | - | - |

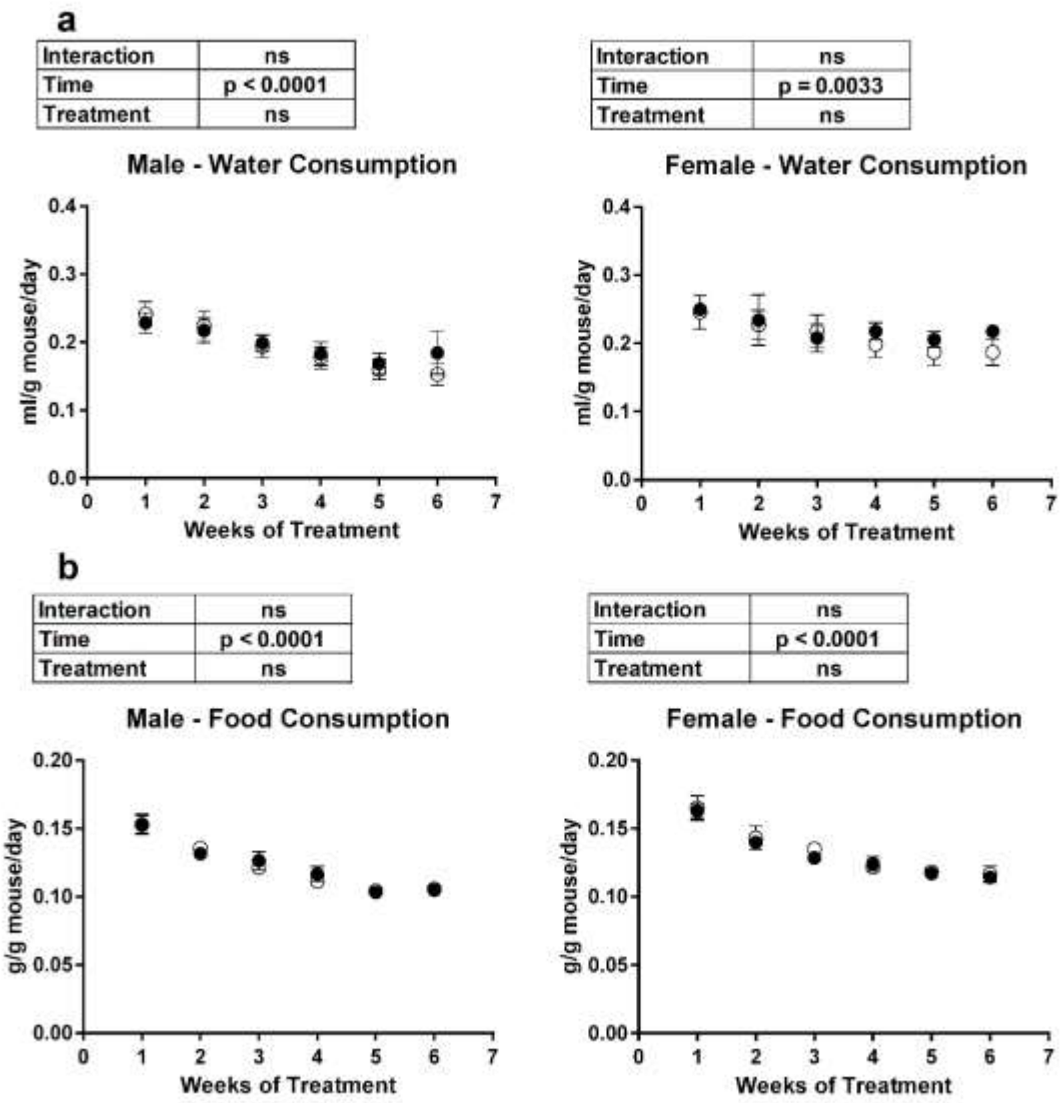

**Fig. S1. Drinking water (a) and food (b) consumption in PFOA-treated mice.**

Three-week-old male and female humanized PPARα and KO mice were treated with either vehicle (Vh, NERL water with 5% sucrose) or PFOA (8 µM in NERL water with 5% sucrose) as drinking water for 6 weeks. During treatment, the mice were fed an American Diet (see Table S1). Food and water consumption were determined on a per cage basis weekly and then divided by the total weight of the mice in the cage and by 7 days to determine the consumption rate per gram of mouse. hPPARα and PPARα null mice were co-housed, therefore the data are not genotype specific. Data are presented as mean ± SE. N = 6-7 cages. Statistical analyses indicated in the boxes are from a 2-Factor Repeated Measures ANOVA.
